# Spatial organization of transcript elongation and splicing kinetics

**DOI:** 10.1101/2021.01.28.428713

**Authors:** Alyssa D. Casill, Adam J. Haimowitz, Brian Kosmyna, Charles C. Query, Kenny Ye, Matthew J. Gamble

**Affiliations:** Department of Molecular Pharmacology, Albert Einstein College of Medicine, Bronx, NY, 10461; Department of Cell Biology, Albert Einstein College of Medicine, Bronx, NY, 10461; Department of Epidemiology and Population Health, Albert Einstein College of Medicine, Bronx, NY, 10461; Department of Systems and Computational Biology, Albert Einstein College of Medicine, Bronx, NY, 10461

## Abstract

The organization of the genome in three-dimensional space has been shown to play an important role in gene expression. Specifically, facets of genomic interaction such as topologically associated domains (TADs) have been shown to regulate transcription by bringing regulatory elements into close proximity^1^. mRNA production is an intricate process with multiple control points including regulation of Pol II elongation and the removal of non-coding sequences via pre-mRNA splicing^2^. The connection between genomic compartments and the kinetics of RNA biogenesis and processing has been largely unexplored. Here, we measure Pol II elongation and splicing kinetics genome-wide using a novel technique that couples nascent RNA-seq with a mathematical model of transcription and co-transcriptional RNA processing. We uncovered multiple layers of spatial organization of these rates: the rate of splicing is coordinated across introns within individual genes, and both elongation and splicing rates are coordinated within TADs, as are alternative splicing outcomes. Overall, our work establishes that the kinetics of transcription and splicing are coordinated by the spatial organization of the genome and suggests that TADs are a major platform for coordination of alternative splicing.

## Introduction

Production of functional messenger RNA (mRNA) is the result of spatial and temporal organization of several highly-regulated processes: RNA Polymerase II (Pol II) recruitment to the promoter, release of Pol II from the promoter-proximal pause, Pol II elongation, splicing, and transcript cleavage and polyadenylation^2^. Elongation and splicing rates are two key kinetic parameters that have been shown to regulate alternative splicing outcomes via the “window of opportunity” model^3–5^. Elongation rates have been measured between 1-6 kb/min (depending on the gene, cell type, and condition)^6–9^. Intron half-lives have been measured between 30 sec and greater than 10 minutes across a variety of systems^10–13^. However, none of the methods to determine splicing kinetics take the wide range of possible elongation rates into account, leading to inaccurate splicing rate measurements. In order to investigate where and how these two kinetic processes work to regulate alternative splicing outcomes, there is a need for an assay that can measure accurate co-transcriptional splicing rates, which requires the concurrent measurement of local elongation rates. Furthermore, while the three-dimensional organization of the genome is known to play a role in regulating transcription, its role in RNA processing, specifically splicing, remains largely unknown. Here we use a novel assay, SKaTER-seq (Splicing Kinetics and Transcript Elongation Rate by Sequencing), to measure rates of transcription and co-transcriptional splicing and ultimately uncover multiple layers of spatial organization of these rates as well as the alternative splicing outcomes they regulate.

## Results

### Accurate estimation of splicing rates requires accurate elongation rates

In order to investigate the contribution of each kinetic process involved in nascent RNA biogenesis (transcription activation, Pol II elongation, splicing, and transcript cleavage) to the final pool of mRNA produced, we built a novel mathematical state model of transcription and co-transcriptional processing. The model describes all possible paths that a region of RNA can go through during its life cycle, representing all intermediates of mRNA biogenesis (Fig. 1A, Extended Data Fig. 1A). The model allows us to calculate the amount of nascent RNA present anywhere within the gene and at any time point following the onset of transcription, incorporating the impact the kinetic parameters influencing nascent RNA production and processing.

**Figure 1.**
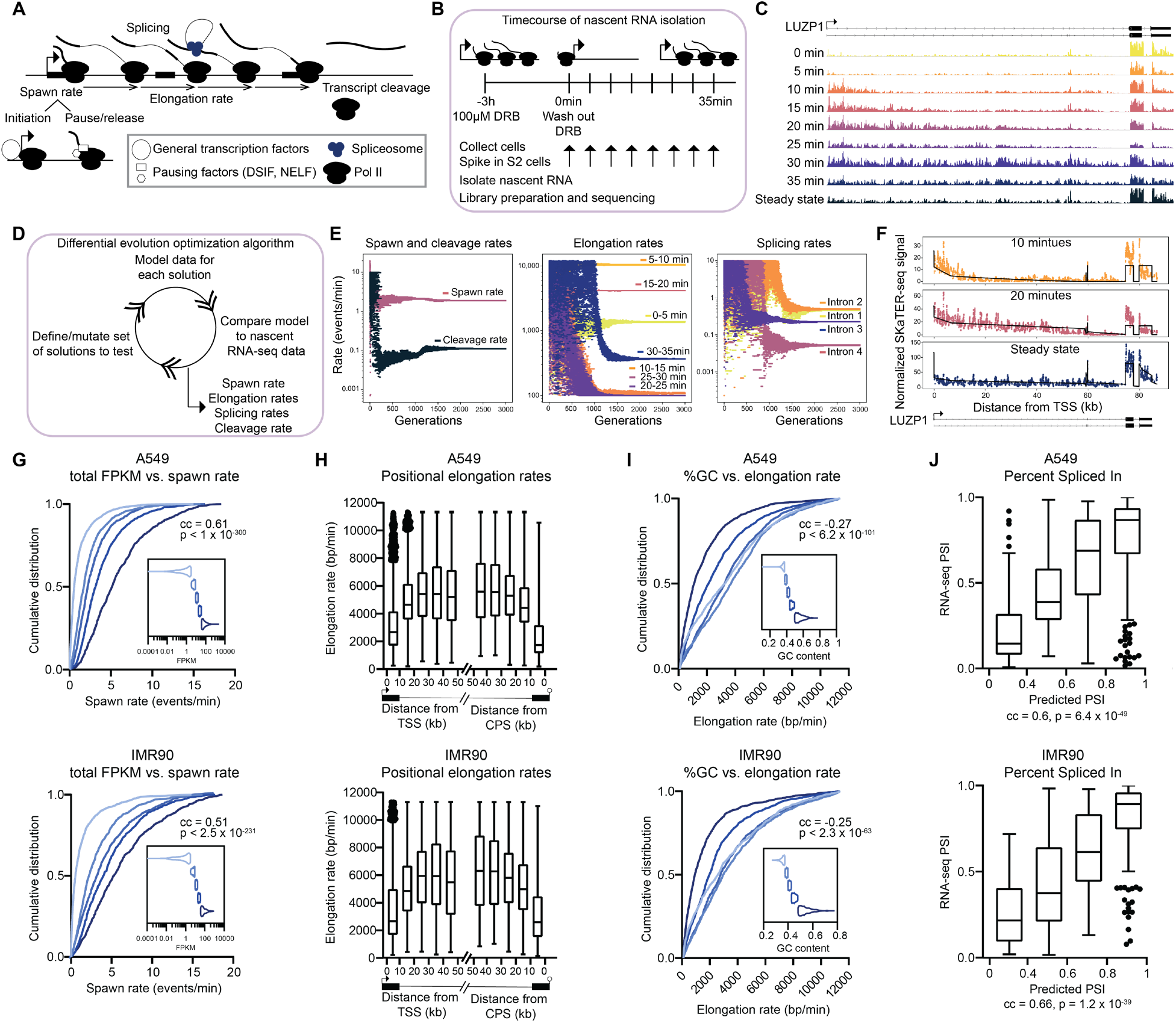
The SKaTER-seq method, computational pipeline, and validation. A. Diagram describing the input parameters (spawn rate, elongation rates, splicing rates, and cleavage rate) for the mathematical model of nascent RNA transcription and co-transcriptional processing. B. Diagram outlining the SKaTER-seq sample preparation procedure. Cells are treated with DRB for 3 hours. DRB is then washed out and cells are collected (with an S2 cell spike-in) every 5 minutes for 35 minutes. Nascent RNA is isolated from each time point and sequenced. C. Example IMR90 SKaTER-seq coverage on the LUZP1 gene across all time points. D. Outline of SKaTER analysis pipeline. The SKaTER-seq analysis pipeline utilizes a differential evolution optimization algorithm coupled with a novel mathematical model of nascent RNA biogenesis and processing to find the rates that best fit the SKaTER-seq data. E. Scatterplots showing the rates of each parameter in the population over the generations of the differential evolution optimization process. F. Example SKaTER-seq coverage tracks (colored lines) on the LUZP1 gene plotted with the modeled coverage from the best fit solution for this gene (black lines). G. Cumulative distribution plots showing the relationship between total RNA levels (FPKM values from RNA-seq) and spawn rates in A549 cells (top) and IMR90 cells (bottom). The genes were sorted by their FPKM values and divided into 5 equal sized bins (the distribution of FPKM values in each bin is represented in the violin plot inset, A549 n = 907 genes/group, IMR90 n = 705 genes/group). The cumulative distribution of the spawn rates of the genes in each bin is plotted on a linear scale. A549 correlation coefficient = 0.61, p < 1 x 10^−300^, IMR90 correlation coefficient = 0.51, p = 2.5 x 10^−231^, calculated using a two-tailed Spearman Rank Correlation test. H. Tukey box plots showing the distribution of the elongation rates within in the denoted window in A549 cells (top) and IMR90 cells (bottom). Center line of boxes represents the median value. N genes per window can be found in Supplemental Table 2. I. Cumulative distribution plots showing the relationship between GC content in a window and the elongation rate in that window in A549 cells (top) and IMR90 cells (bottom). The windows were sorted by their GC content and divided into 5 equal sized bins (the distribution of the GC content in each bin is represented in the violin plot inset, A549 n = 1216 windows/group, IMR90 n = 901 windows/group). The cumulative distribution of the elongation rates of the windows in each bin is plotted on a linear scale. A549 correlation coefficient = −0.27, p = 6.2 x 10^−101^, IMR90 correlation coefficient = −0.25, p = 2.3 x 10^−63^, calculated using a two-tailed Spearman Rank Correlation test. J. Tukey box plots comparing the predicted Percent Spliced In (PSI) based on SKaTER-seq rates to PSI measured using RNA-seq data in A549 (top) and IMR90 cells (bottom). Predicted PSI was modeled using the SKaTER-seq rates for genes containing alternative splicing events. P values, as indicated, calculated using a two-tailed Spearman Rank Correlation test. Center line of boxes represents the median value. Additional statistical summary information can be found in Supplemental Table 2.

Previous studies on the timing of intron removal do not incorporate local elongation rate measurements into their determination of splicing rates^12–14^. While assuming an average elongation rate is likely sufficient when studying organisms with relatively short genes (*S. cerevisiae, S. pombe, D. melanogaster*), we hypothesized that to determine accurate splicing rates of introns in relatively long mammalian genes, incorporation of local elongation rates would be crucial. By keeping exon-exon junctional coverage constant in our model, we were able to determine the relationship between elongation rate and apparent splicing rate on a model gene with the 3’SS of an intron 30 kb from the cleavage site. We found that a linear change in elongation rate results in an exponential change in apparent splicing rate (Extended Data Fig. 1B), demonstrating that concurrent evaluation of splicing and elongation rates is necessary. SKaTER-seq is the first assay to measure locus-specific elongation rates in order to accurately determine co-transcriptional splicing rates.

### SKaTER-seq measures kinetics of nascent RNA production, Pol II elongation, splicing, and transcript cleavage

SKaTER-seq measures rates of transcription activation, Pol II elongation, splicing, and transcript cleavage genome-wide in mammalian cells from nascent RNA-sequencing. Sequencing is performed at several timepoints following transcriptional block and release using 5,6-Dichlorobenzimidazole 1-β-D-ribofuranoside (DRB), a reversible small molecule inhibitor of P-TEFb (positive transcription elongation factor b)^15^ (Fig. 1B, methods). The resulting SKaTER-seq read coverage shows progression of transcription into the body of genes following DRB washout (Fig. 1C).

Read coverage from SKaTER-seq at any position in a gene is a function of the various rates that contribute to nascent RNA biogenesis and processing. While it is difficult to measure these rates directly from nascent RNA-seq coverage, we can describe the contributions of each rate to the coverage using our mathematical state model of transcription and co-transcriptional processing. To find rates that optimally fit the measured nascent RNA-seq coverage over the time course, we coupled our model with an evolutionary optimization algorithm^16,17^ (Fig. 1D). For each gene, spawn rate (which incorporates both Pol II initiation rate and promoter-proximal pause release rate), elongation rate (approximated as a constant-time process and evaluated in up to seven “speed-zones,” one per timepoint), splicing rate of each intron, and transcript cleavage rate are determined concurrently and iteratively improved by the optimization algorithm until the solution converges on the best fit (Fig. 1E). The optimal solution contains estimated rates for each parameter describing a single gene, which can be used to recapitulate the coverage pattern in SKaTER-seq data (Fig. 1F).

### SKaTER-seq rates of transcription and RNA processing are validated by orthogonal biological data

We performed SKaTER-seq in two biological replicates of both IMR90 lung fibroblasts and A549 lung adenocarcinoma cells. Within each biological replicate of each cell line, our algorithm was able to measure rates on ~50% of expressed genes. (Extended Data Fig. 2A, methods). We examined the various rates impacting transcription and splicing outcomes genome wide, and found that the spawn, elongation, splicing, and cleavage rates are highly consistent across biological replicates. Additionally, each rate type exhibited a wide distribution within both cell lines (Extended Data Fig. 2A-C, Supplemental Table 1).

To validate the SKaTER-seq rates, we compared our data to orthogonal biological data. For example, steady state mRNA levels are controlled by the rate of production of new mRNA molecules (spawn rate) and the rate of RNA decay of each transcript. Therefore, spawn rates should correlate with (but not perfectly match) steady state mRNA levels. We performed total RNA-seq in both cell lines and, consistent with expectations, found strong correlation between spawn rate and steady state mRNA levels (Fig. 1G). In contrast, cleavage rates should not impact RNA production or decay. Consistently with this, we see only a weak correlation between cleavage rates and steady state mRNA abundance (Extended Data Fig. 3A).

Consistent with previous reports^6,7,18^, we find that elongation rates tend to accelerate over gene bodies and decelerate at the end of genes (Fig. 1H). We also confirm that elongation rates are negatively correlated with GC content^7,8^ (Fig. 1I).

Our data shows that the median nascent intron half-life in A549 cells is ~3 minutes, while in IMR90 the median half-life is ~1.5 minutes, consistent with previous observations that mammalian splicing occurs on the order of minutes^19^. It has been previously reported that the minor spliceosome (an orthogonal complex that removes specific introns harboring distinct splice site consensus sequences^20^) removes introns slower than the major spliceosome^10^. This is consistent with our data in A549 cells, where minor spliceosome introns^21^ are removed slower than major spliceosome introns. However, in IMR90 cells, there is not a significant difference between the two types of introns (Extended Data Fig. 2B). Using RT-qPCR for the U2 and U12 small nuclear RNAs (snRNAs) that work analogously in major and minor spliceosomes, respectively, we found that the relative level of U12 expression was significantly elevated in IMR90 cells, which potentially explains why minor spliceosome introns are removed more efficiently in these cells (Extended Data Fig. 2B).

Finally, we used our model, including the splicing and elongation rates impacting cassette exon alternative splicing events, to predict the percent of transcripts that contain a cassette exon at steady state (PSI, percent spliced-in) and found a significant positive correlation with actual PSI values determined from total RNA-seq data (Fig 1J). The fact that our predicted PSI values are consistent with measured PSI values validates our model and our estimates of splicing rates. Overall, these results show that SKaTER-seq produces highly reproducible and biologically meaningful kinetics for nascent RNA biogenesis and processing genome-wide.

### Splicing rates are coordinated within gene bodies

The spliceosome must assemble *de novo* on each intron to complete splicing catalysis. Therefore, intron removal is often thought to be an autonomous process, as each intron harbors the cis-acting elements necessary for its own removal. While crosstalk between introns has been shown to promote recruitment of spliceosomes^22,23^, it is unclear if there is any relationship between the splicing rates of introns within the same gene. To examine this, we compared the rate of a constitutive intron to the rate of its adjacent downstream constitutive intron and found significant correlation between these groups in both A549 and IMR90 cells (Fig. 2A-B). Indeed, splicing rates of constitutive introns appear to be clustered within genes (Fig. 2C-D). In order to quantify the extent and significance of this gene-level clustering, we calculated the variance of the log_2_ transformed rates within each gene, then the variance of all genes was summed to determine the total observed variance. We then permuted the splicing rates genome-wide and recalculated the total variance 100,000 times in order to obtain the expected distribution of the variance under the null hypothesis that splicing rates are not coordinated within genes (Fig. 2E). We found that the observed total variance is significantly less than would be expected by random chance, indicating that there is a high degree of intragenic coordination among introns (Fig. 2F-G).

**Figure 2.**
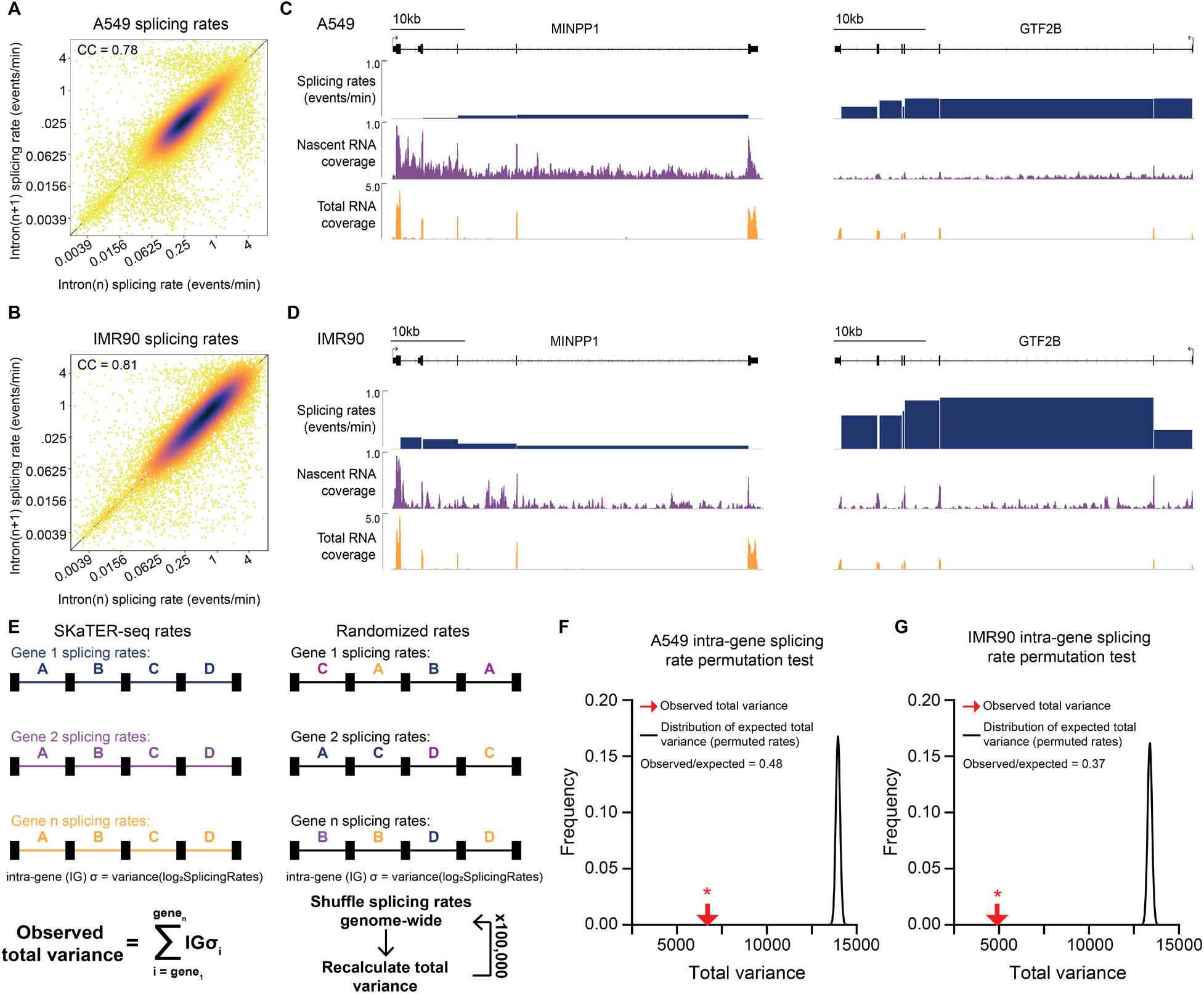
Splicing rates are coordinated at the gene level. A-B. Scatterplot representing the relationship between the rate of splicing of an intron (intron(n)) and the rate of splicing of the intron immediately downstream (intron(n+1)) in A549 cells (24,349 pairs of introns, correlation coefficient = 0.78, p < 1 x 10^−300^, A) and IMR90 cells (16,938 pairs of introns, correlation coefficient = 0.81, p < 1 x 10^−300^, B). Correlation coefficients and p values calculated using a two-tailed Spearman Rank Correlation test. Only constitutive introns were included in this analysis. Gradient from blue to yellow indicates decreasing density of points (kernel density estimation). C-D. Example genome browser tracks showing the splicing rate of each intron (blue), the nascent RNA-seq coverage (purple), and the total RNA-seq coverage (orange) in A549 cells (C) and IMR90 cells (D). E. Overview of the variance metric and permutation test used to determine the level of coordination of intra-gene splicing rates. F-G. Comparison of the observed variance (red arrow) and the distribution of the total variance of the permuted rates (black line) across 100,000 permutations in A549 cells, (F) and IMR90 cells (G). * p < 1 x 10^−5^, representing the number of times (out of 100,000) the total variance based on randomized data was less than the observed total variance. A549 = 33,922 introns, IMR90 = 24,262 introns included in the permutation analysis.

### Splicing and elongation rates are coordinated within gene neighborhoods

To determine if splicing rates are coordinated beyond gene boundaries, we performed a distance-filtered pairwise correlation of introns. In this analysis, we observe that the correlation of splicing rates persists to distances greater than the length of most genes (Extended Data Fig. 4A). To extend our analysis beyond gene boundaries, we performed a similar distance-filtered pairwise correlation of gene-average rates (to eliminate any bias of intragenic coordination) and found that the autocorrelation of elongation rates and splicing rates persists through extended distances (Fig. 3A). To determine the optimal size of gene neighborhoods exhibiting this coordination, we split the genome into equally sized windows (at increasing window sizes) and performed a similar permutation test as described above, but instead calculated the total intra-window variance of intron-level splicing rates (Extended Data Fig. 4B). We found that the ratio between the observed total variance and expected total variance (O/E) is significantly below 1 in windows up to 10 Mb long, indicating that intronic splicing rates are highly coordinated even within large genomic regions (Extended Data Fig. 4C). Furthermore, using gene-average rates and the same coordination test in genomic windows of increasing size, we found that gene-average splicing and elongation rates, but not spawn and cleavage rates, are significantly coordinated within large genomic regions (Extended Data Fig. 4D).

**Figure 3.**
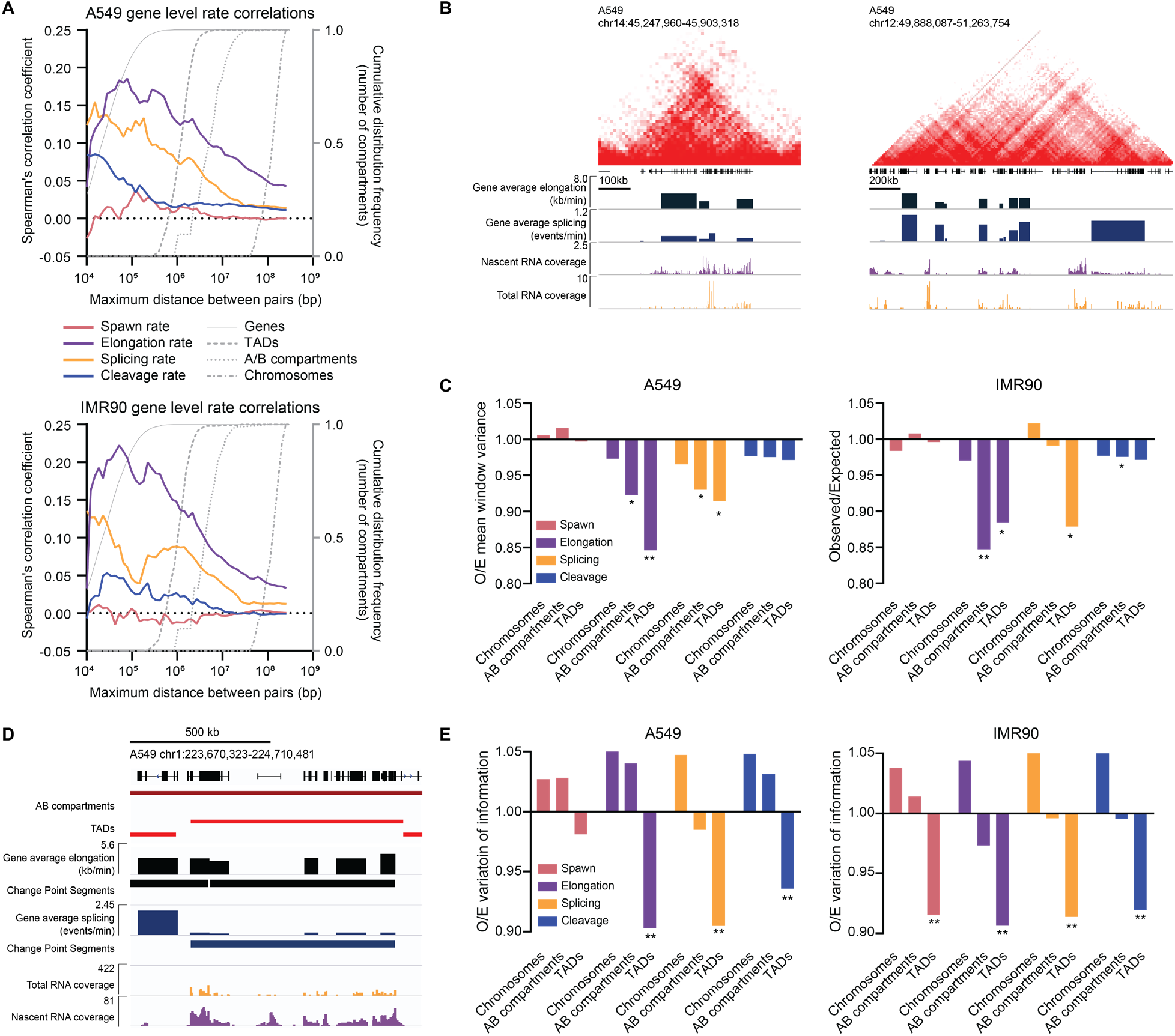
Splicing rates and elongation rates are coordinated within high-order chromatin domains. A. Two-tailed spearman rank correlation of gene average splicing rates for paired genes filtered for the indicated maximum distance. Grey lines (right y-axis) represent cumulative distributions for the lengths of each type of genomic compartment at the distances marked on the x-axis. B. Example genome browser tracks from A549 Hi-C data **and** SKaTER-seq data showing coordination of gene-average splicing rates (dark blue) and elongation rates (blue), along with the nascent RNA-seq coverage (purple) and the total RNA-seq coverage (orange) within TADs. C. Observed/expected mean window rate variance in A549 (left) and IMR90 cells (right) for the indicated rates. ** p < 1×10^−5^, * p < 0.01, from a one-tailed z-test comparing the observed variance to the distribution formed by the permutations. D. Example genome browser tracks from A549 Hi-C data (Encode Project Consortium, 2012) and SKaTER-seq data showing the genomic segments and change points based on the SKaTER-seq rates compared to AB compartments and TADs. E. Observed/expected variation of information in A549 (left) and IMR90 cells (right) for the indicated rates in the indicated compartments. ** p < 1×10^−5^ from a one-tailed z-test comparing the observed variation of information to the distribution formed by the permutations. Additional statistical summary information can be found in Supplemental Table 2.

### Splicing and elongation rates are organized within higher-order chromatin domains

The window sizes wherein splicing and elongation rates are coordinated are reminiscent of well-established genomic compartments defined by the three-dimensional organization of chromatin. Topologically associated domains (TADs) are large genomic regions (100 kb – 2 Mb) that have a high frequency of interaction and function to keep genes and regulatory elements in close spatial proximity^24^. A and B compartments are larger domains that separate transcriptionally active and inactive chromatin^25^. The location and distribution of A/B compartments and TADs can be determined using a technique known as Hi-C, which uses high-throughput sequencing to determine local chromatin interactions^25^. Using published Hi-C data from IMR90 and A549 cells, we defined A/B compartments and TADs in a unified fashion using previously published analysis tools^24,26,27^ (methods). The previously noted persistence of gene-average splicing and elongation rate pairwise correlation extends well into the range of TAD and A/B compartment sizes (Fig. 3A). Indeed, we see evidence that SKaTER-seq rates within TADs tend to have similar gene-average splicing rates and elongation rates (Fig. 3B, Extended Data Fig. 5A). Using TADs, A/B compartments, or chromosomes as the genomic windows in the coordination test described above (Extended Data Fig. 4B), we find that elongation rates are significantly coordinated within TADs and A/B compartments. We also find that splicing rates are significantly coordinated within TADs in both cell lines and A/B compartments in A549 cells (Fig. 3C). Overall, this demonstrates that elongation and splicing kinetics are organized within regions of enriched local chromatin interaction.

This compartment-based coordination analysis shows that TADs and A/B compartments segment elongation and splicing rates in a statistically significant fashion. We then took an unsupervised approach using a binary segmentation change point detection algorithm to segment the data based only on the SKaTER-seq rates, and compared those segments to TADs, A/B compartments, and chromosomes (Extended Data Fig. 5B-C, Fig. 3D). We found that for all rate types (except spawn rates in A549 cells), the optimal groupings of genes based on their rates is significantly more similar to TADs than would be expected by random chance, and do not significantly resemble A/B compartments or chromosomes (Fig. 3E). Taken together, these results demonstrate that TAD boundaries segment the genome in a way that organizes genes to coordinate their transcription elongation and splicing rates.

### Changes in elongation-rate sensitive alternative splicing patterns are coordinated within TADs

It has been shown that elongation rates can influence alternative splicing outcomes by regulating the window of time a splice site has to be recognized and used before a competing event occurs^5^ (Fig. 4A). Whether or not an exon is included or excluded during this “window of opportunity” is also dependent on the splicing rates. We found that both elongation rates and splicing rates are coordinated within TADs (Fig. 3). Given that elongation and splicing kinetics are two key processes that govern alternative splicing outcomes, these observations led to the dramatic prediction that changes in alternative splicing patterns should also be coordinated at the TAD level (Fig. 4B). To test this, we used a coordination test similar to the tests described above. We quantified the changes in exon inclusion (changes in PSI, or ΔPSI) between two RNA-seq datasets, then converted the ΔPSI to either +1 or −1 based on the direction of change to get the ordinal ΔPSI, which was used to calculate the total intra-window variance. We then shuffled the ΔPSI values and recalculated the total ordinal ΔPSI intra-window variance over 100,000 trials to determine the expected distribution under the null hypothesis. (Extended Data Fig. 6A).

**Figure 4.**
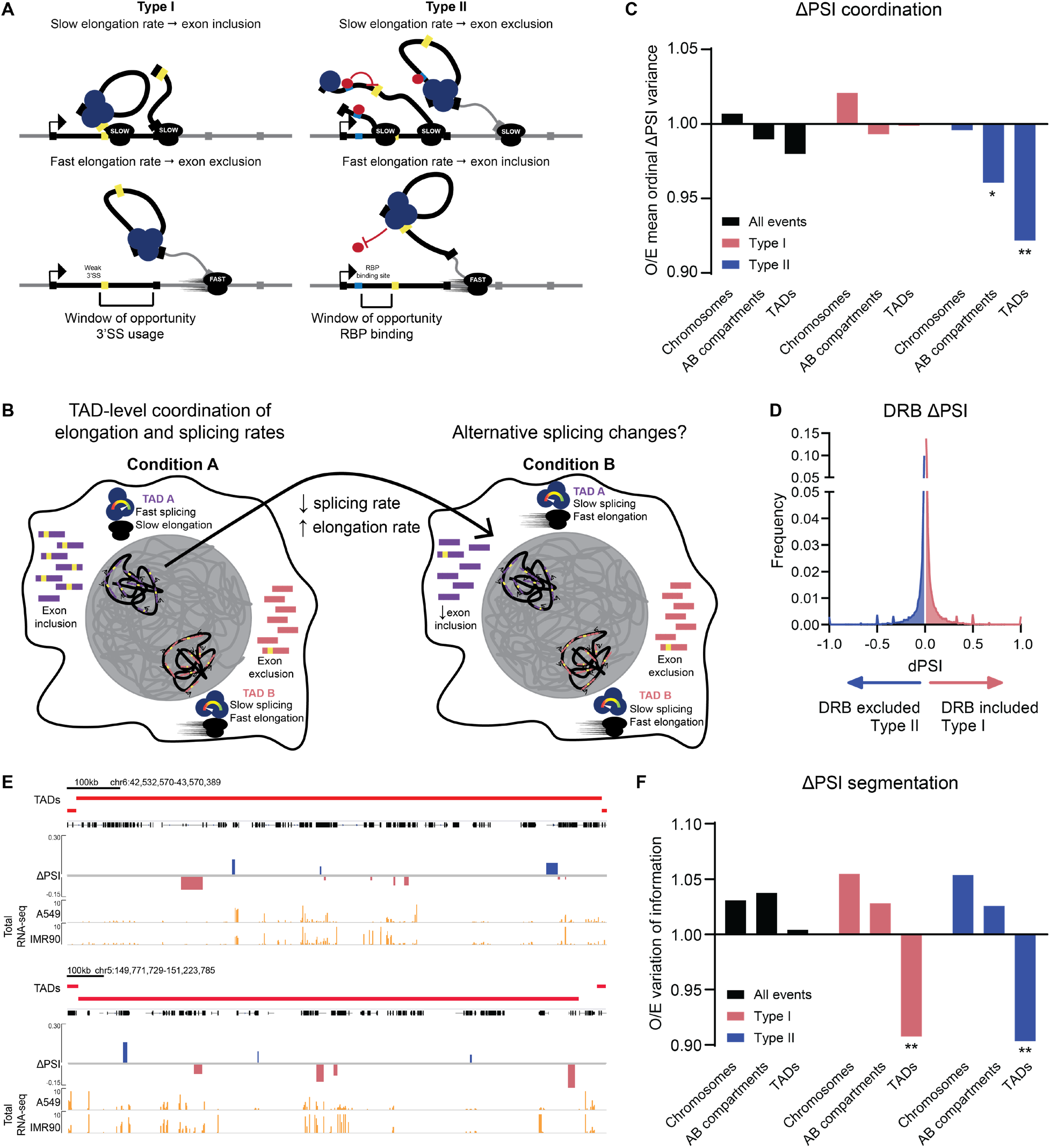
Elongation rate sensitive alternative splicing outcomes are coordinated within higher-order chromatin domains. A. Schematic of type I and type II alternative splicing events. Type I cassette exons increase in PSI as elongation rates decrease. Type II cassette exons increase in PSI as elongation rates increase. B. Proposed model wherein changes in coordinated splicing and elongation rates within TADs are accompanied by coordinated changes in alternative splicing patterns. C. Observed/expected ordinal ΔPSI variance metric for all events, type I events, and type II events in the indicated compartment types (based on A549 Hi-C data). ** p < .001, * p < 0.05, from a one-tailed z-test comparing the observed ordinal ΔPSI variance to the distribution formed by the permutations. D. Distribution of percent spliced in (PSI) levels for each alternative splicing event (n = 81,199) in A549 cells treated with 25 μM DRB for 24 hours. Events that are more included in the treated condition are enriched for type I events, and events that are more excluded in the treated condition are enriched for type II events. E. Example genome browser tracks from A549 Hi-C data and SKaTER-seq data showing coordination of changes in PSI between IMR90 and A549 cells along with the total RNA-seq coverage (orange) within TADs. Type I events are in pink and type II events are in blue, positive values indicate more inclusion in IMR90 cells and negative values indicate more inclusion in A549 cells. F. Observed/expected variation of information for all events, type I events, and type II events in the indicated compartments (based on A549 Hi-C data). ** p < 1×10^−5^ from a one-tailed z-test comparing the observed variation of information to the distribution formed by the permutations. Additional statistical summary information can be found in Supplemental Table 2.

Using RNA-seq data from A549 and IMR90 cells, we found that the differences in the alternative splicing seen between these cell lines are not significantly coordinated within the compartment types tested (Fig. 4C). However, there are two main and opposing mechanisms by which elongation rate influences alternative splicing outcomes, previously defined as type I and type II events (Fig. 4A)^5^. In type I events, decreased elongation rate causes an increased probability of exon inclusion, as it gives additional time for a weak 3’SS in the upstream intron to be recognized and used. In type II events, decreased elongation rate causes a decreased probability of exon inclusion, as it gives additional time for negative regulators to bind in the upstream intron and prevent 3’SS recognition. Therefore, a compartment-level change in elongation would have opposite effects on these distinct types of events. To analyze these events separately, we defined a list of elongation rate-sensitive events using total RNA-seq from A549 cells treated with 25μM DRB, which has been previously shown to decrease Pol II elongation rates^28^. Events with increased inclusion in DRB treated cells are enriched for type I, and events with decreased inclusion enriched for type II (Fig. 4D). We observed that type I and type II events tend to have opposite ΔPSI directions within TADs, and their direction is clustered based on their event type (Fig. 4E). In fact, we found that type II events are significantly coordinated within both TADs and A/B compartments (Fig. 4C). The lack of coordination of type I events is due to the fact that these events seem to be globally coordinated, as the majority of the type I ΔPSI values are negative (more included in A549 cells, Extended Data Fig. 6B). Furthermore, we found that unsupervised clustering of both types of events produces genomic segments that are significantly more similar to TADs than would be expected by random chance (Fig. 4F). Taken together, these results uncover a previously unappreciated level of coordination of alternative splicing dependent on the three-dimensional organization of the genome, implying a broad mechanism of regulation of exon inclusion coordination.

## Discussion

SKaTER-seq is a novel genome-wide method that measures the kinetics of nascent RNA biogenesis and processing, including Pol II elongation rates and splicing rates. There are several assays that have been used in the past to measure elongation rates (e.g. GRO-seq, BruDRB-seq^6,8^) and, similarly, there are several techniques used to track the life-cycle of an intron (RT-qPCR, single molecule imaging, metabolic labeling, long-read nascent RNA-seq^10–13,29,30^). However, SKaTER-seq is the first assay to measure these two parameters simultaneously genome-wide and to incorporate locus-specific elongation rates into the measurement of splicing rates. This is a crucial step forward, as accurate determination of co-transcriptional splicing rates requires accurate measurements of elongation rates at that locus.

Previous studies have shown that the splice sites of adjacent introns can act synergistically to more effectively recruit spliceosomes^22,23^, which could also facilitate the stabilization of rates within genes. Metabolic labeling studies in *D. melanogaster* showed that introns within the same gene have consistent half-lives^12^. Analysis of our measured rates shows that co-transcriptional splicing kinetics are highly coordinated within gene bodies in human cells (Fig. 2). This is potentially a result of regulation from the promoter (promoter strengths have been previously shown to regulate splice site selection^31^) or cross-talk among spliceosomes on adjacent introns as described in the model of exon definition^22^. Alternatively, this coordination could be driven by introns within a gene that evolved to have similar spliceosome or splicing factor recruitment efficiency or sequences that promote similar catalytic efficiency of the spliceosome.

We found that elongation and splicing rates as well as elongation-rate sensitive alternative spicing outcomes are coordinated within TADs, suggesting that these are areas of local kinetic regulation of gene expression. It is possible that by bringing genes into close proximity, TADs can coordinate alternative splicing outcomes on a multi-gene scale. While further studies are necessary to determine the mechanism of TAD-level coordination of splicing and transcription, it is possible that TADs can function as zones of splicing and elongation factor enrichment. Splicing factors, particularly SR proteins, have been shown to form condensates in the cell^32^, and extensive research has gone into understanding the contents and functions of these condensates. It has recently been shown that the phosphorylation status of Pol II can influence the types of condensates that it associates with during transcription (hyperphosphorylated Pol II CTD associates with splicing factor rich condensates, whereas hypophosphorylated Pol II associates with mediator condensates)^33^. We suggest that the proximity of a TAD to a splicing factor or elongation factor rich condensate may drive a TAD-level effect on transcription and splicing kinetics, and ultimately regulate the direction of alternative splicing within a TAD. The potential for a gene to regulate the alternative splicing outcomes in neighboring genes by concentrating transacting factors is a novel regulatory mechanism that may have a widespread impact on cellular function. Overall, our work evokes many new and exciting questions about how the organization of transcription and RNA processing kinetics is controlled and how it contributes to gene expression outcomes.

## Acknowledgments

We thank G.A. Hamilton and V. Gupta for their bioinformatic help. We thank the Einstein Genomics Core Facility for experimental support and the Einstein High Performance Computing Core for computer cluster administration. Supported by R01CA155232 and R01GM134279 (M.J.G), R01GM57829 (C.C.Q), P30CA013330 (Albert Einstein College of Medicine Cancer Center Support core grant), and NIH training grant 5T32GM007491-41 (A.D.C and A.J.H.).

## Author contributions

Conceptualization: A.D.C., M.J.G.; Methodology: A.D.C., K.Y., M.J.G.; Software: A.D.C., M.J.G.; Formal Analysis: A.D.C., K.Y.; Investigation: A.D.C., A.J.H., B.K.; Writing – original draft preparation: A.D.C.; Writing – review and editing: A.D.C., A.J.H., B.K., C.C.Q., K.Y., M.J.G; Supervision: M.J.G., C.C.Q.

## Data and materials availability

All sequencing data produced for this study has been deposited at the GEO database (www.ncbi.nlm.nih.gov/geo/), accession number GSE146290. Published Hi-C data used in this study can be found at GSE63525 (IMR90) and GSE105600 (A549). The SKaTER-seq analysis software is available at https://github.com/TheRealGambleLab.

---

**Extended Data Figure 1.**
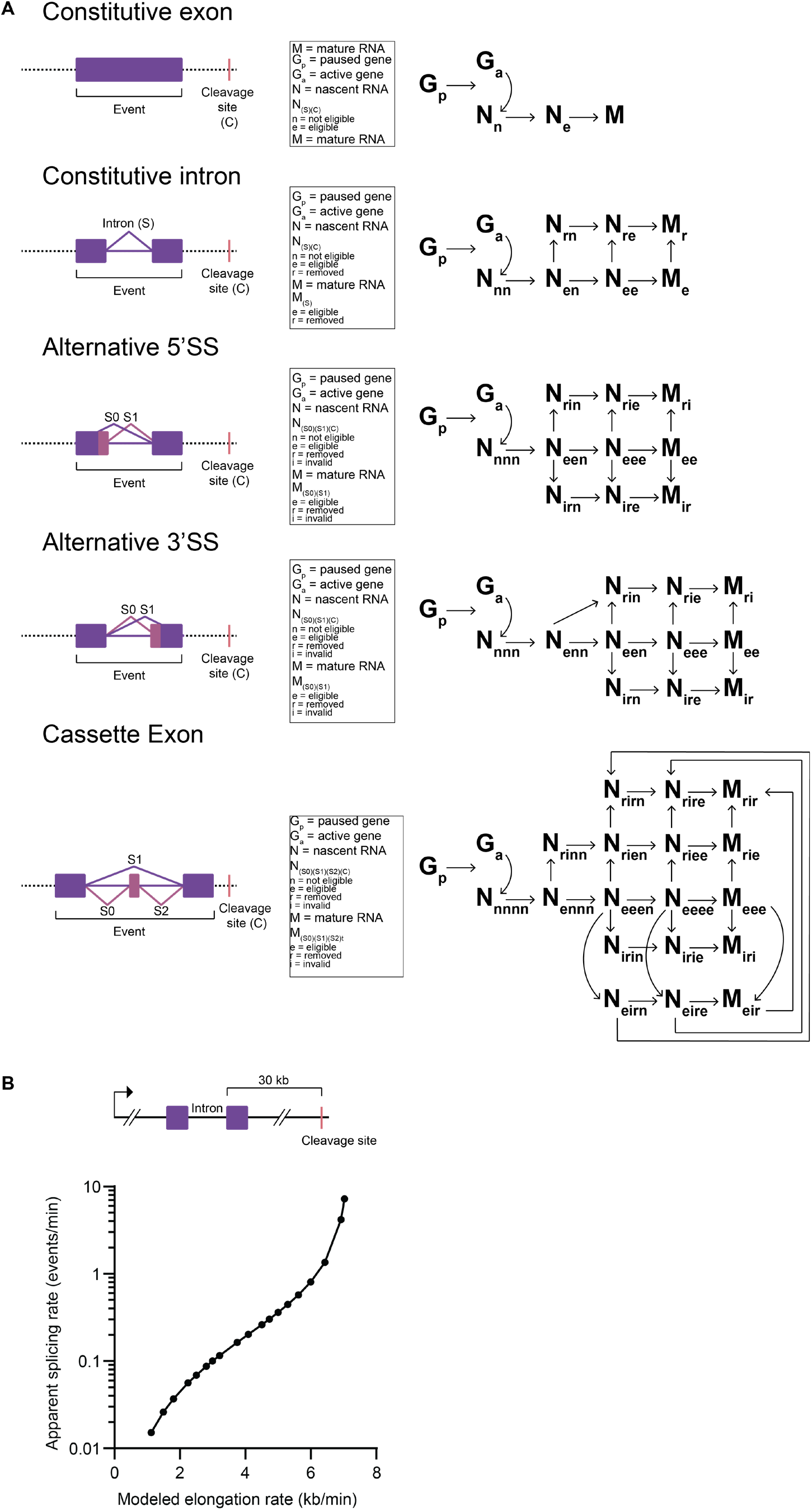
SKaTER-seq state models. A. State models for constitutive exons, constitutive introns, alternative 5’SS, alternative 3’SS, and cassette exons. B. Modeling the effect of different elongation rate on apparent splicing rate given a constant junctional coverage. Top: Diagram of conceptual gene used to model junctional coverage at different rates. Bottom: Relationship between modeled elongation rate and apparent splicing rate. The data demonstrates that for an intron with a 3’SS 30 kb from the cleavage site, a linear change in elongation rate is met with an exponential change in apparent splicing rate.

**Extended Data Figure 2.**
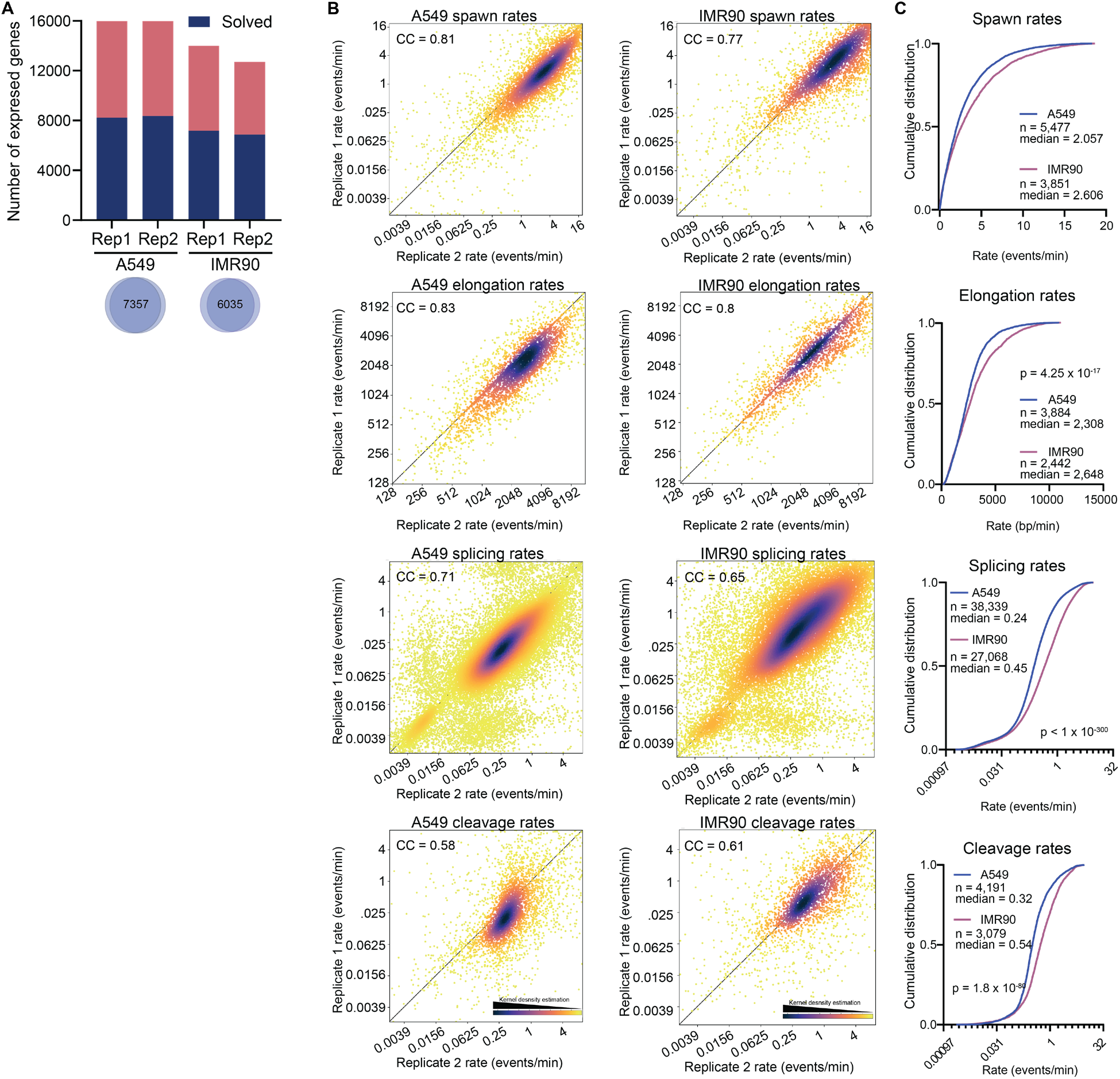
Summary of rates measured using SKaTER-seq. A. Bar graph showing the total number of genes expressed in each replicate (passing the cutoff of 5 reads/kb), and the fraction of those genes solved by the SKaTER-seq algorithm. Venn diagrams below the graph show the number of genes that are solved in both biological replicates within each cell line. B. Scatterplots showing the correlation of each rate (spawn, elongation, splicing and cleavage) across the two biological replicates in each cell line. A549 spawn rate correlation coefficient (CC) = 0.81, p < 1 x 10^−300^, IMR90 spawn rate CC = 0.77, p < 1 x 10^−300^, A549 cleavage rate CC = 0.58, p < 1 x 10^−300^, IMR90 cleavage rate CC = 0.61, p = 2.04 x 10^−318^, A549 elongation rate CC = 0.83, p < 1 x 10^−300^, IMR90 elongation rate CC = 0.87, p < 1 x 10^−300^, A549 splicing rate CC = 0.71, p < 1 x 10^−300^, IMR90 splicing rate CC = 0.65, p < 1 x 10^−300^. Correlation coefficients are calculated using a two-tailed Spearman Rank Correlation test. C. Cumulative distribution plots showing the spawn, elongation, splicing, and cleavage rates in IMR90 and A549 cells. P values, as indicated, calculated using a two-sided Kolmogorov-Smirnov (KS) test. Additional statistical summary information can be found in Supplemental Table 2.

**Extended Data Figure 3.**
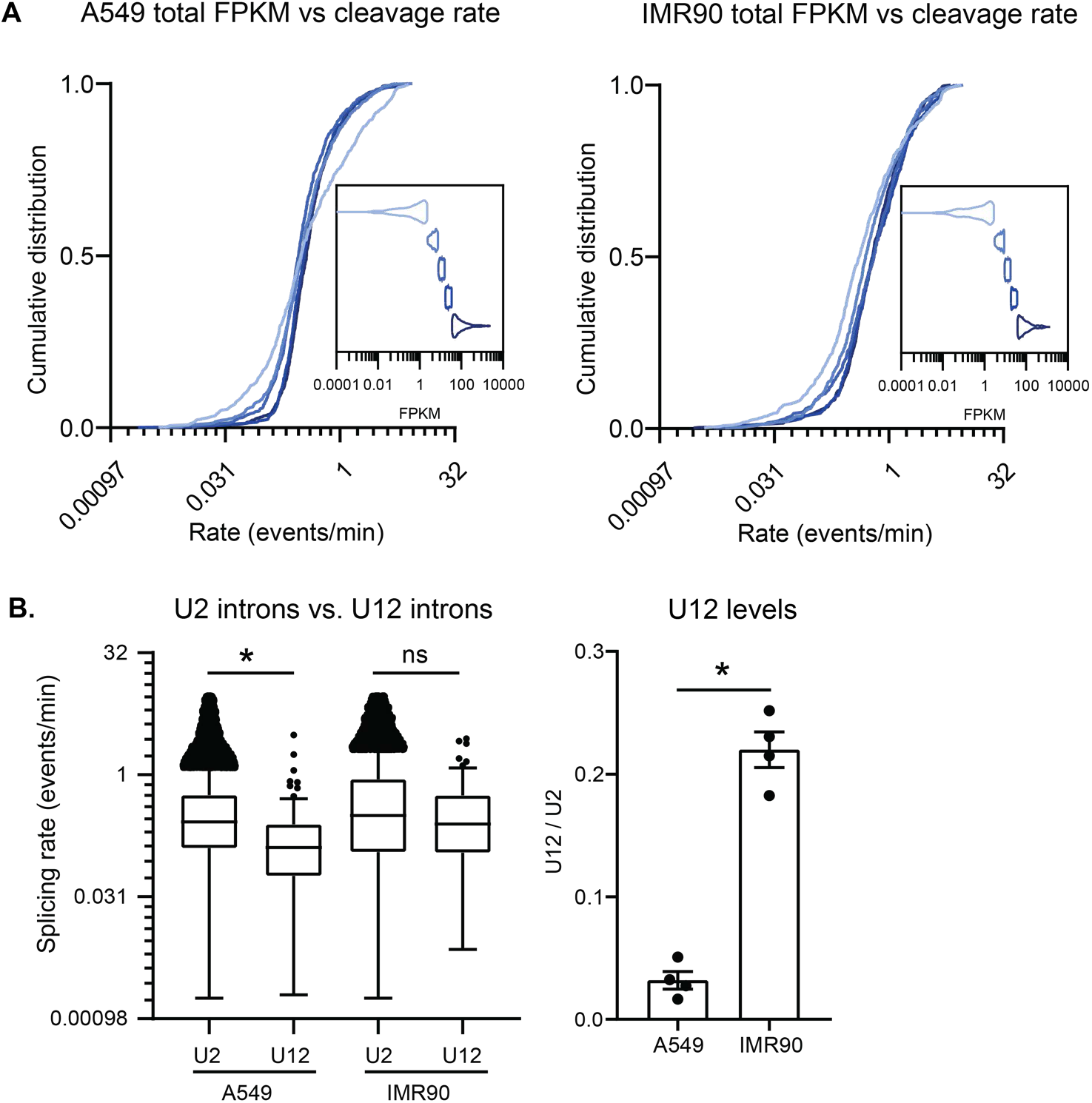
Additional rate validation using orthogonal data. A. Cumulative distribution plots showing the relationship between total RNA levels as determined using FPKM values from RNA-seq and cleavage rates in A549 cells (left) and IMR90 cells (right). The genes were sorted by their FPKM values and divided into 5 equal sized bins (the distribution of FPKM values in each bin is represented in the violin plot inset, A549 n = 685 genes/group, IMR90 n = 562 genes/group). The cumulative distribution of the cleavage rates of the genes in each bin is plotted on a log_2_ scale. A549 CC = 0.06, p = 3 x 10^−4^, IMR90 CC = 0.12, p = 2.3 x 10^−11^, Correlation coefficients are calculated using a two-tailed Spearman Rank Correlation test. B. Left: Splicing rates of introns removed by the major and minor spliceosome in A549 and IMR90 cells. A549 n U2 introns = 38,277, n U12 introns = 62, IMR90 n U2 introns = 27,024, n U12 introns = 44, * p = 1.2 x 10^−5^ calculated using a two-sided Mann-Whitney U test to compare the log of the rates between groups (A549 Mann-Whitney U statistic = 819212, IMR90 Mann-Whitney U statistic = 534351). Center line of boxes represents the median value. Right: Ratio of U12 to U2 expression in A549 and IMR90 cells. Error bars represent SEM of four biological replicates. * p = 0.028, calculated using a two-sided Mann-Whitney U test (Mann-Whitney U statistic = 0) to compare the log of the rates between groups.

**Extended Data Figure 4.**
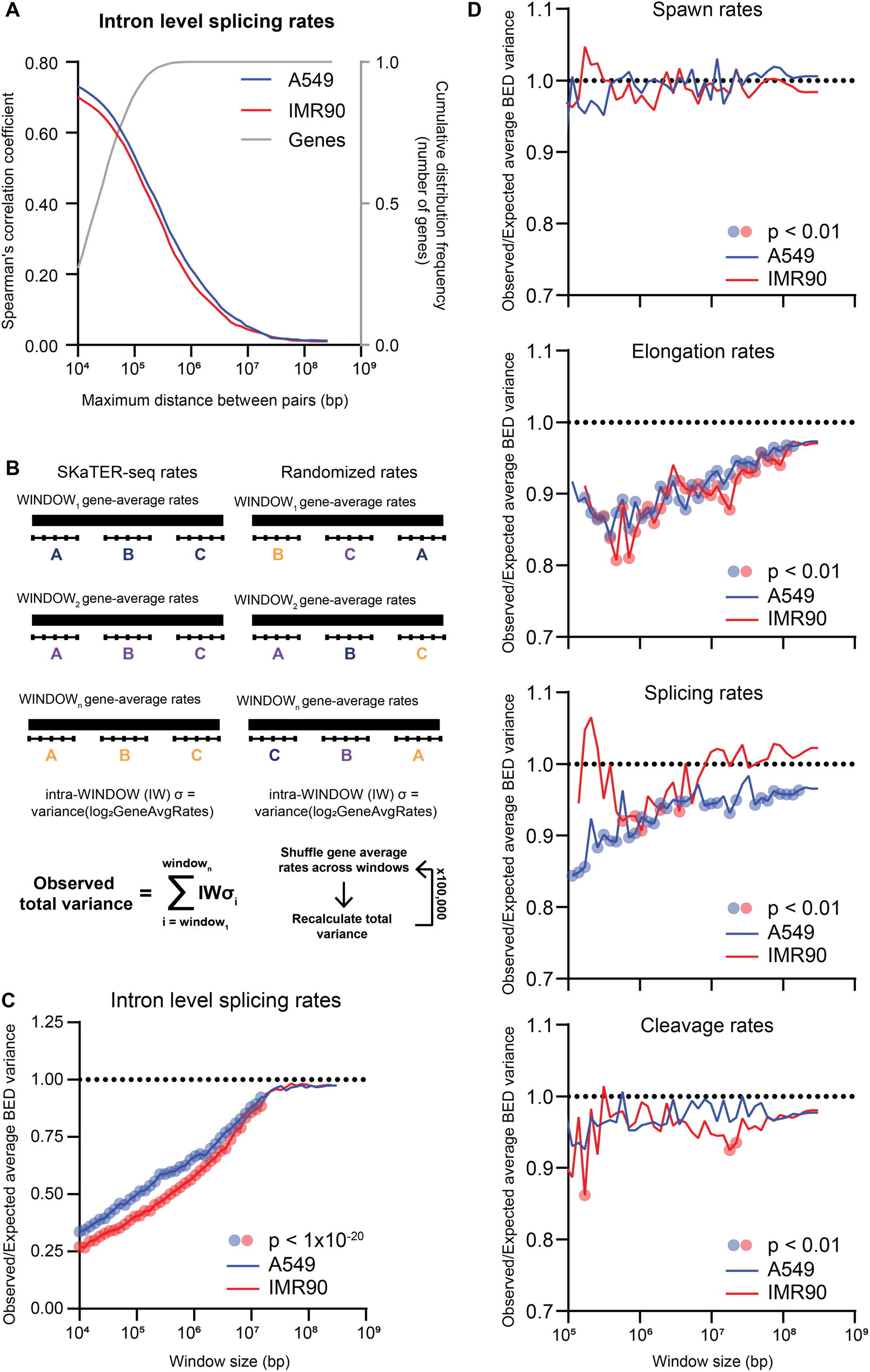
Correlation and coordination of rates extend into local gene neighborhoods. A. Two-tailed spearman rank correlation of splicing rates for paired introns filtered for the indicated maximum distance. Grey lines (right y-axis) represent cumulative distribution for the gene lengths covered at the distances marked on the x-axis. B. Overview of the variance metric and permutation test used to determine the level of coordination of intra-window rates. C. Using the permutation test described in B, where variance is calculated based on all introns within the window. For each window length (x-axis), the genome was divided into windows of equal length (per chromosome), the observed variance and expected variance were calculated, and the ratio of the two (observed/expected) is plotted on the y-axis. Windows significant p-values (as indicated on graph) from a one-tailed z-test comparing the observed variance to the distribution formed by the permutations are marked with translucent circles. D. Using the permutation test described in B, where the variance is calculated based on the gene-average rates. For each window length (x-axis), the genome was divided into windows of equal length (per chromosome), the observed variance and expected variance were calculated, and the ratio of the two (observed/expected) is plotted on the y-axis. Windows significant p-values (as indicated on graph) from a one-tailed z-test comparing the observed variance to the distribution formed by the permutations are marked with translucent circles. Additional statistical summary information can be found in Supplemental Table 2.

**Extended Data Figure 5.**
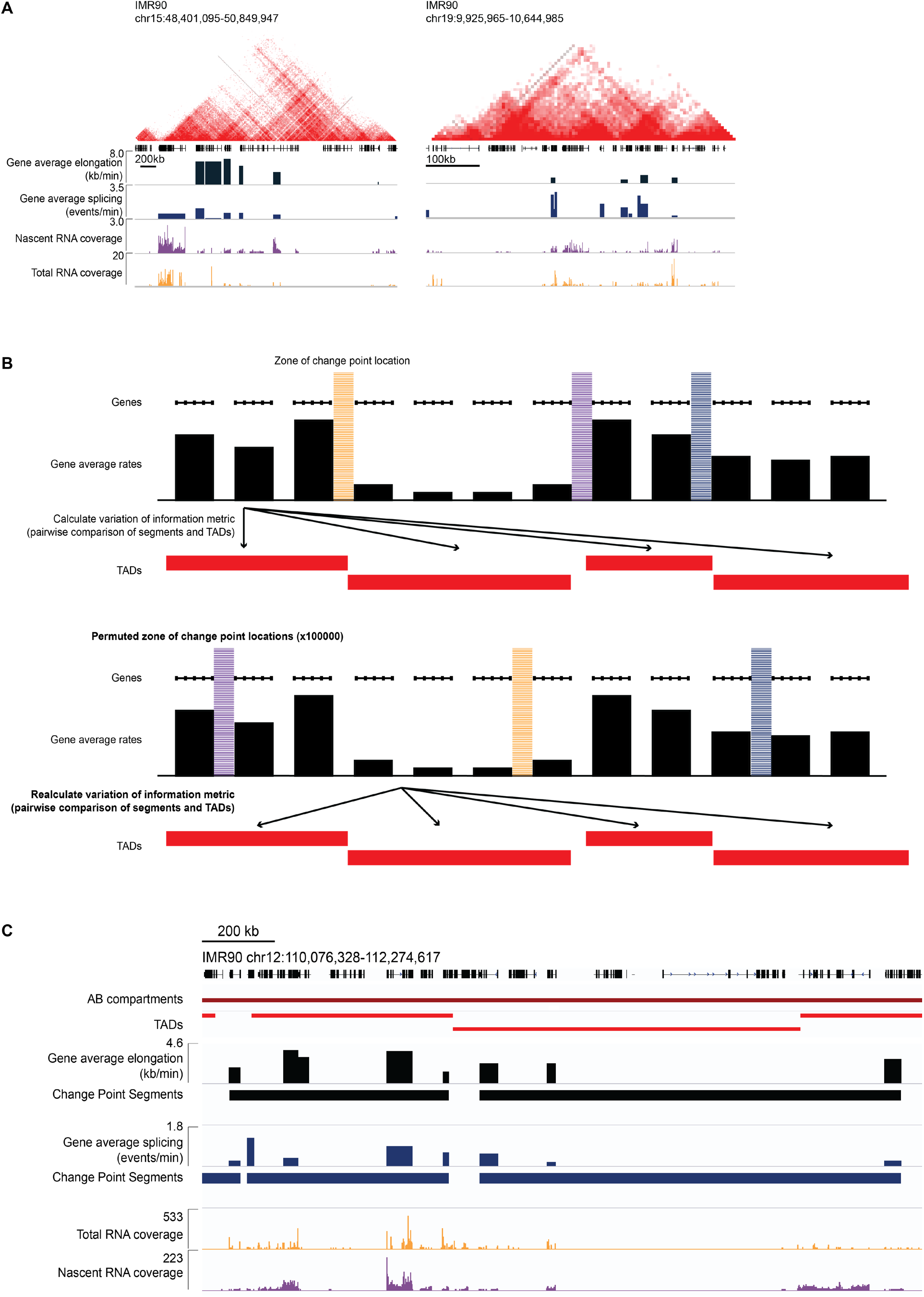
Genome segmentation and TADs. A. Example genome browser tracks from IMR90 Hi-C data and SKaTER-seq data showing coordination of gene-average splicing rates (dark blue) and elongation rates (blue), along with the nascent RNA-seq coverage (purple) and the total RNA-seq coverage (orange) within TADs. B. Schematic of the change point detection program and the permutation test used to compare rate-based change points with compartment boundaries. C. Example genome browser tracks from IMR90 Hi-C data and SKaTER-seq data showing the genomic segments and change points based on the SKaTER-seq rates compared to AB compartments and TADs.

**Extended Data Figure 6.**
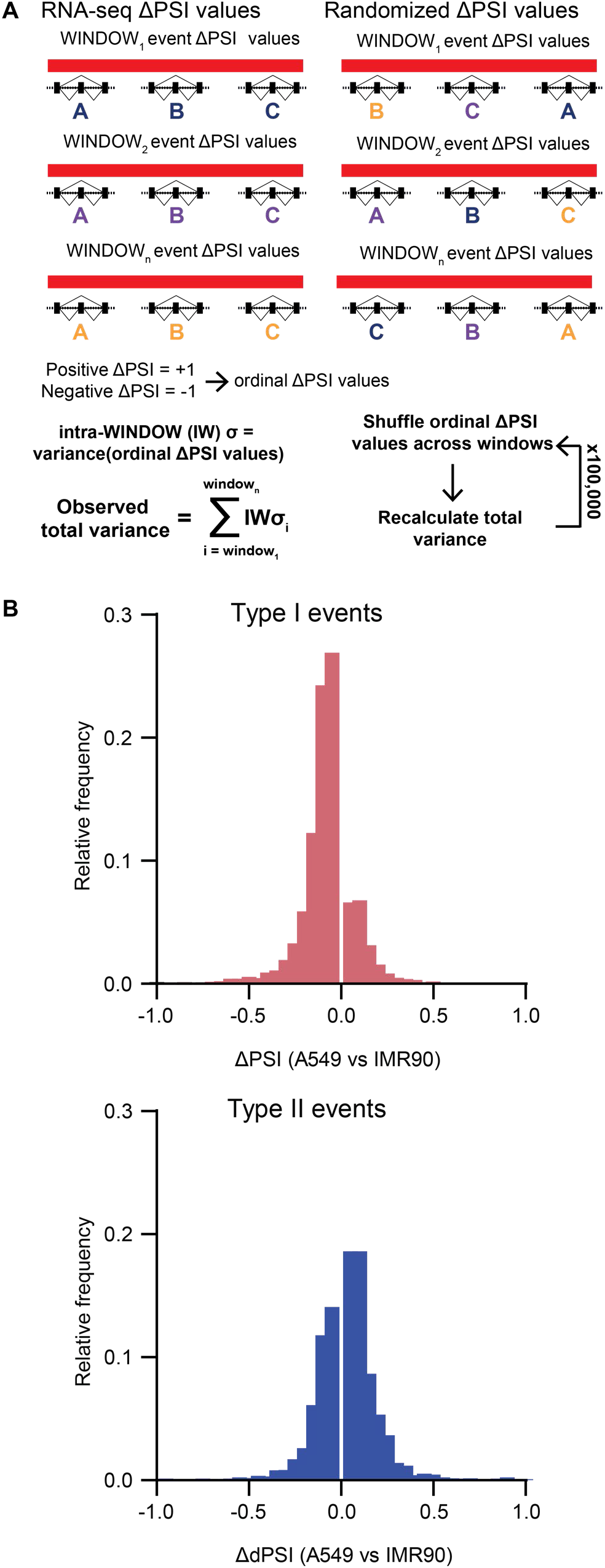
Coordination of alternative splicing outcomes. A. Overview of the ordinal ΔPSI variance metric and permutation test used to determine the level of coordination of alternative splicing outcomes. B. Histograms of the distribution of ΔPSI values (comparing PSI in A549 and IMR90 cells) for previously established type I (n = 4707) or type II (n = 3094) events. Positive values indicate more inclusion in IMR90 cells and negative values indicate more inclusion in A549 cells.

**Supplemental Table 1. SKaTER-seq spawn, elongation, splicing, and cleavage rates.**

Each tab contains the named SKaTER-seq rates for both biological replicates within a cell line.

The columns are as follows:

Gene name, location (TSS for spawn rates, intron start/intron stop for splicing rates, no location for elongation rates, CPS for cleavage rates), rate in biological replicate 1, rate in biological replicate 2, average rate.

**Supplemental Table 2. Statistics Summary**

Each tab contains the statistical information relevant for the named figure panel.

## Materials and Methods

### Cell lines and culture conditions

IMR90 primary human fetal lung fibroblast cells (ATCC CCL-186) were cultured in MEM supplemented with 10% fetal bovine serum (FBS) and 1x Penicillin-Streptomycin (pen-strep) solution. A549 lung cancer cells (ATCC CCL-185) were cultured in RPMI supplemented with 10% FBS and 1x pen-strep solution. Drosophila melanogaster S2 cells (ATCC CRL-1963) were cultured at 28C in Schneider's Drosophila Medium supplemented with 10% heat-inactivated FBS and 1x pen-strep solution.

### Total RNA purification and RNA-sequencing

Cells were grown to 90% confluence in 6 cm dishes. Where noted, cells were treated with with 25 μM 5,6-Dichlorobenzimidazole 1-β-D-ribofuranoside (DRB, Sigma D1916-50MG) or DMSO (Fisher Scientific BP231100) as a vehicle control for 24 hours. RNA was isolated using TRIzol (Invitrogen, 15596018) according to the manufacturer’s protocol. mRNA was enriched using oligo(dT) beads and sequenced to a target depth of 40 million reads per sample (150bp paired-end reads, non-stranded mRNA-seq) by Novogene.

### qPCR for snRNAs

RNA was isolated as described above. cDNA from 500 ng of total RNA was reverse transcribed using Invitrogen Superscript III reverse transcriptase with 50 pmol of random hexamer primers in a 20 ul reaction, according to manufacturer’s protocol. Relative snRNA levels were determined by RT-qPCR. snRNA qPCR primers and diluted cDNA were added to Applied Biosystems™ Power™ SYBR™ Green Master Mix for 10 ul reaction in 384 well plate. qPCR was performed on Applied Biosystems ViiA 7 Real Time PCR system. After 10 minutes of incubation at 95 °C, 40 cycles of amplification (95 °C for 15 s, 60 °C for 1 minute) were measured and melting point analysis was performed to ensure specificity. The efficiency-corrected threshold cycle (ΔCT) method was used to determine the relative levels of snRNA. Total U2 snRNA is the sum of RNU2-1 and RNU2-2P. Primer sequences are listed below.

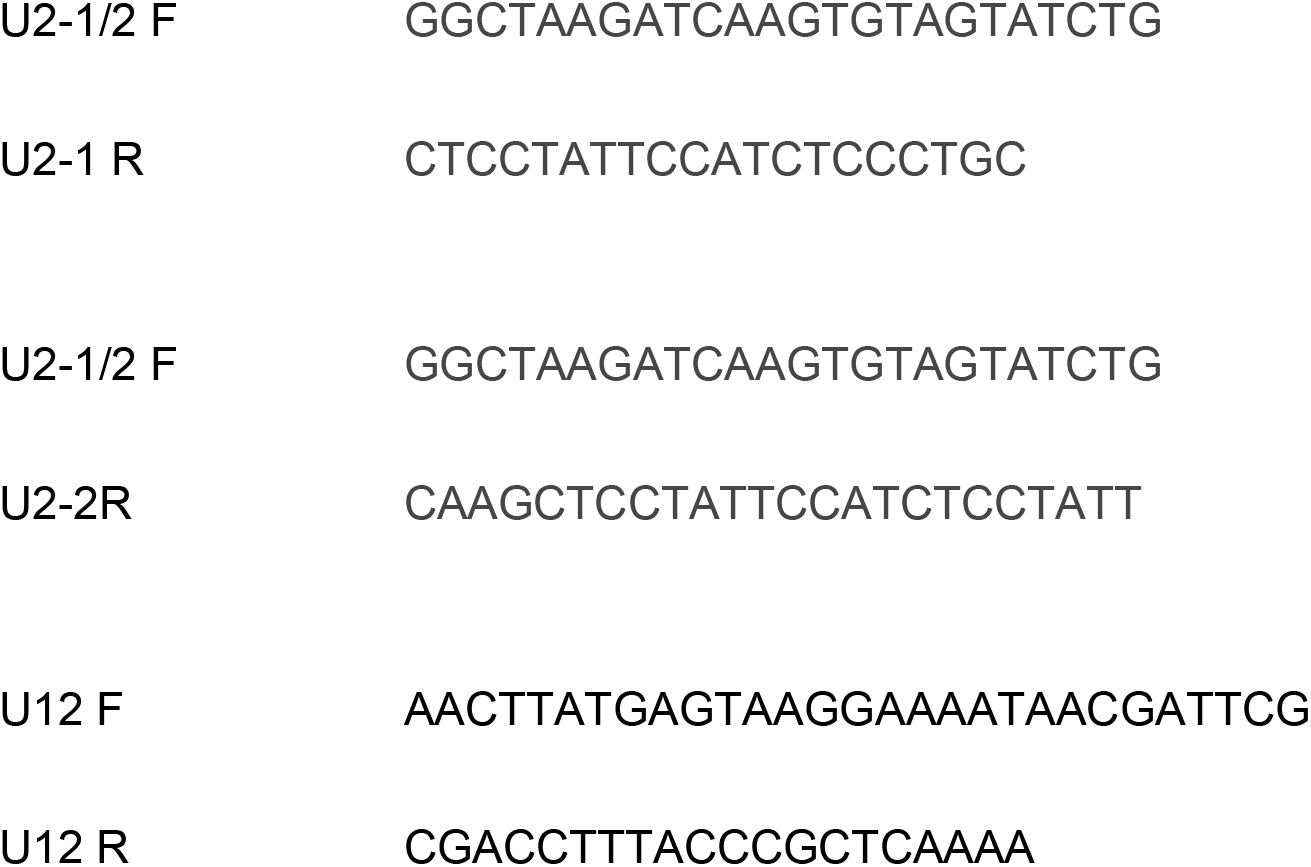

### Total RNA-seq analysis

Reads were aligned to the reference human genome (hg19) using STAR (v2.6.1b^34^) with outFilterMultimapNmax = 1. MarkDuplicates (picard-tools v2.3.0^35^) was used to filter out PCR duplications from the mapped reads using the default parameters. Cufflinks (v2.2.1^36^) was used to measure the Fragments Per Kilobase of transcript per Million mapped reads (FPKM) using the default parameters. PSI values were calculated using rMATS (v3.2.5^37^) unpaired analyses and the default parameters.

### SKaTER-seq

#### Treatment and time course

Our cell treatment and nascent RNA isolation protocol is loosely adapted from several previous studies and has been optimized for our method^10,14,38–41^. Cells were grown to 90% confluence in nine 10 cm dishes (one dish per time point). Cells were treated with 100 μM 5,6-Dichlorobenzimidazole 1-β-D-ribofuranoside (DRB, Sigma D1916-50MG) or DMSO (Fisher Scientific BP231100) as a vehicle control for 3 hours. After 3 hours, cells were washed twice in 37 °C 1x PBS and 37 °C fresh media was added to each plate, except the 0-minute time point, which was never washed out. Cells were immediately put back in the 37 °C incubator. At each 5 minute time point, one plate of cells was removed from the incubator, quickly washed once in ice cold 1x PBS, then 1mL of Cytoplasmic Lysis Buffer + Drosophila S2 cells (CL+S2 buffer, 25 mM Tris pH 7.9, 150 mM NaCl, 0.1 mM EDTA, 0.1% TritonX, 1 mM DTT, 1x protease inhibitor cocktail, 1.3×10^6^ S2 cells/mL) was added directly to the plate. Cells were collected by scraping and kept on ice until the end of the time course.

#### Nascent RNA Isolation

After all samples were collected and incubated in CL+S2 buffer on ice for at least 10 min, the lysates were centrifugated at 3000 RPM for 5 min at 4 °C. The supernatant was removed and the pellet was washed in 1 mL CL buffer (without S2 cells) and re-centrifugated at the same speed. The pellet was then resuspended in 100 μL Glycerol Resuspension Buffer (GR buffer, 20 mM Tris pH 7.9, 75 mM NaCl, 0.5 mM EDTA, 50% glycerol, 0.85 mM DTT), to which 1.1 mL Nuclear Lysis Buffer (NL buffer, 20 mM HEPES pH 7.6, 300 mM NaCl, 7.5 mM MgCl2, 1% NP-40, 1 mM DTT, 1 M Urea) was subsequently added. The resulting lysate was incubated on ice (with regular vortexing) for 15 min, then centrifugated at 16,000 G for 10 min at 4 °C. The resulting chromatin pellet was resuspended in TRIzol (Invitrogen, 15596018) and RNA was extracted according to the manufacturer’s protocol. To further deplete mRNA, the NEBNext Poly(A) mRNA Magnetic Isolation Module (NEB, E7490L) was used to capture and remove polyA+ mRNA. The manufacturer’s protocol was followed until the mRNA was bound to the beads. Following mRNA binding, the supernatant was removed from the beads and the remaining nascent RNA was ethanol precipitated and resuspended in ddH_2_O.

#### Nascent RNA library preparation and sequencing

Nascent RNA was quantified on a Denovix QFX fluorometer using the Qubit RNA HS assay according to the manufacturer’s protocol. The quality of the RNA was assessed using an Agilent 2100 Expert Bioanalyzer instrument. Sequencing libraries were generated using the KAPA Stranded RNA-seq Kit with RiboErase (HMR) (Roche, 07962282001) according to the manufacturers protocol. Final libraries were also subjected to bioanalyzer runs to ensure proper fragment length distribution. All 9 libraries were barcoded and sequenced together on two lanes of an Illumina HiSeq (paired-end 150 bp reads, stranded total RNA-seq).

#### Mapping nascent RNA-seq reads

RNA-seq reads were aligned using STAR (v2.6.1b^34^) to a custom genome containing both the human (hg19) and drosophila (dm3) genomes using the following parameters: outFilterMultimapNmax = 1, outFilterScoreMinOverLread = 0.51,outFilterMatchNminOverLread = 0.51, outFilterMismatchNmax = 4, alignIntronMax = 50000, twopassMode = basic.

MarkDuplicates (picard-tools v2.3.0^35^) was used with the default parameters to mark PCR duplications within the mapped reads.

### SKaTER-seq rate calling

Parameters used during the SKaTER-seq analysis pipeline are described below and summarized in Table 1.

**Table 1.** Parameters used in the SKaTER-seq analysis pipeline.

|  | Parameter | Value used | Description |
| --- | --- | --- | --- |
| Experimental parameters | Timepoints | 0, 5, 10, 15, 20, 25, 30, 35, steady state | Time of collection (post-DRB washout) for each sample |
| Read filtering criteria | Step | 10 | Evaluate every $n^{\text{th}}$ base pair from read coverage and during modeling |
|  | Fraction masked | 0.5 | In the “local variance filter,” this fraction of reads will be removed based on their variance across time points |
|  | TSS filter data index | 1 | Index of the sample used to calculate TSS usage |
|  | TSS merger distance | 150 bp | Distance below which multiple annotated start sites will be merged |
|  | Cleavage site merger distance | 150 bp | Distance below which multiple annotated cleavage sites will be merged |
| Coverage normalization | Betas ($\beta$) | See methods and table 2 | Drosophila coverage within timepoint sample / Drosophila coverage in steady state sample, for each sample (steady state $\beta = 1$) |
| Rate bounds | DRB release rate lower bound | 0.00166 events/min | Upper and lower limits for each rate type – rates will be optimized within these bounds. |
|  | DRB release rate upper bound | 10.0 events/min |  |
|  | Spawn rate lower bound | 0.00166 events/min |  |
|  | Spawn rate upper bound | 20 events/min |  |
|  | Elongation rate lower bound | 100 bp/min |  |
|  | Elongation rate upper bound | 12,000 bp/min |  |
|  | Splicing rate lower bound | 0.00166 events/min |  |
|  | Splicing rate upper bound | 10 events/min |  |
|  | Cleavage rate lower bound | 0.00166 events/min |  |
|  | Cleavage rate upper bound | 10 events/min |  |
| Contamination estimation parameters | $\gamma_t$ | See methods and table 3 | Drosophila exonic coverage / Drosophila intronic coverage for each sample |
| | Alpha ($\alpha$) upper bound | Minimum $\gamma_t$ value | Lowest $\gamma_t$ based on observed drosophila exonic and intronic read counts |
| | Alpha ($\alpha$) lower bound | 0.31 | Theoretical lower limit based on the drosophila genome if splicing never occurred |
| Optimization parameters | Population size (p) | 100 | Parameters of the differential evolution optimization algorithm |
|  | Mutation scale factor (f) | 0.6 |  |
|  | Crossover probability (Cr) | 0.8 |  |
|  | Max generations | 10,000 | Stop optimization if rates have not converged |
|  | Critical (maxRSS-minRSS) | 0.1 | If the (maxRSS – minRSS) is below this for a parameter, the optimization has converged on a single answer. |

#### Read filtering

Nascent RNA-seq has additional challenges that must be addressed before analyzing the data. First, because our library preparation protocol does not include a de-capping step, we do not generally get coverage within the first ~100 nt of a gene. To keep this consistent across genes, we mask the first 100 nt of each gene before starting the analysis. Additionally, we limit our analysis to every 10^th^ nucleotide to increase computational speed.

Introns and other non-coding regions can be highly repetitive, which can prevent accurate read mapping. Furthermore, introns tend to be structured, which can be problematic for the reverse transcription step during library preparation. It is necessary to filter out regions like these across the SKaTER-seq time points without losing valuable information about the changes in nascent RNA coverage over time. In order to achieve this, we developed a “local variance filter” that uses a 1 kb moving window. We calculate the variance across the time points for each nucleotide in the window, then determine if the variance of the central nucleotide is in the top half of the variances of nucleotides in that window. If it is not in the top half of variances, that nucleotide is masked.

#### Data normalization using spike-in

Depending on the time since DRB wash out a sample was collected, we expect samples to have different total levels of nascent RNA. These expected global changes make normalization to total human read counts an inappropriate strategy. Our cross-species spike-in of the same number of untreated (steady state) Drosophila S2 cells to every sample allows us to normalize across time points. Reads that map to the Drosophila genome are used to calculate a scaling factor used to normalize the human data. We define this scaling factor, β, as:

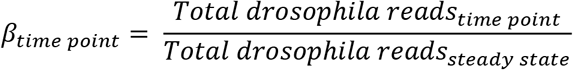

The steady state scaling factor (β_ss_) is set as 1. See Table 2 for the β value for each time point within each biological replicate.

**Table 2.** Betas (β) for normalization of coverage across time points.

| | $\beta_0$ | $\beta_5$ | $\beta_{10}$ | $\beta_{15}$ | $\beta_{20}$ | $\beta_{25}$ | $\beta_{30}$ | $\beta_{35}$ | $\beta_{ss}$ |
| --- | --- | --- | --- | --- | --- | --- | --- | --- | --- |
| IMR90 replicate 1 | 0.462228 | 0.642719 | 0.675147 | 0.815167 | 0.491228 | 0.710548 | 0.683576 | 0.761057 | 1 |
| IMR90 replicate 2 | 0.710914 | 1.110914 | 0.588317 | 0.660694 | 0.75145 | 1.246555 | 0.680906 | 1.153516 | 1 |
| A549 replicate 1 | 0.612234 | 0.398373 | 0.444541 | 0.970889 | 2.008554 | 0.806803 | 1.520069 | 0.552527 | 1 |
| A549 replicate 2 | 0.395925 | 0.290262 | 0.177994 | 0.312852 | 0.592945 | 0.757501 | 0.652494 | 0.898514 | 1 |

#### Gene and event identification

The first step of the SKaTER-seq analysis pipeline is to identify if a gene can be accommodated by our computational pipeline and, if so, what events make up the gene architecture. We started with RefSeq annotated transcription start sites (TSSs) and cleavage and polyadenylation sites (CPS). The current version of our pipeline can only analyze genes with one TSS, as transcription from multiple TSSs confounds the measurement of Pol II elongation rates. If a gene has two or more annotated TSSs, we first calculate the distance between the sites. We consider TSSs within 150 bp of each other to be sufficiently close to be used as a single TSS annotated at the upstream position. This is because our method lacks the resolution to be able to differentiate between usage of start sites that are <150 bp apart. If the start sites are >150 bp apart, we use the coverage at each TSS in the 5-minute sample to determine usage. We calculate the sum over the coverage over the 200 bp downstream of the masked region (described above) and determine the maximum coverage of an annotated TSS for that gene. If any other annotated TSS has greater than 10% of the maximum coverage, the gene is marked as having “multiple start sites” and will not be further analyzed by our algorithm. Otherwise, the TSS corresponding to the window of maximal coverage is considered the dominant TSS and is included in the analysis. On the other end of the gene, we cannot use coverage to determine CPS usage, due to transcriptional readthrough past the annotated end of the gene. Therefore, if a gene has two annotated CPS locations greater than 150 bp apart, the gene is marked as having “multiple cleavage and polyadenylation sites” and is not analyzed further.

If a gene passes the TSS and CPS filters, we then break down the gene into individual events. The possible events are constitutive exons, constitutive introns, alternative 5’ SS usage, alternative 3’ SS usage, cassette exon events, and double cassette exon events (Fig. S1A). If a gene contains any event not included in this list, it will not be analyzed further as we do not currently have the mathematical models needed to model these genes. We use the junctional reads from a gene to identify the introns in a gene, and if any intron overlaps another, we determine the type of alternative splicing that is used within that event. In order to be called an intron, we require that a junction has at least 10% the level of coverage of the maximally covered junction in the gene and that the intron is at least 50 bp long. This eliminates any indels or other sequencing errors from being inaccurately identified as introns.

#### Modeling nascent RNA transcription and co-transcriptional processing

We model the kinetics of nascent RNA through a set of differential equations, based on the assumptions that the elongation rate is deterministic for any given gene, and the time after passing a splice site to completion of splicing and the time after passing a cleavage site to transcript cleavage are both random variables following an exponential distribution. For each event type, we divide total nascent RNA into different states, based on status of the intron, the cleavage event, and transcription progress (Fig. S1A). The change in nascent RNA between states is described by a differential equation. For example, for a constitutive intron in a gene with only one cleavage site, we describe the state of the nascent RNA as N_xy_, with the first subscript, x, representing the status of the intron (not eligible for splicing, n, eligible for splicing, e, or removed, r) and the second subscript, y, representing the status of the cleavage event (not eligible for cleavage, n, or eligible for cleavage, e):

1. N_nn_: nascent RNAs that have not passed the 3’SS, which are not eligible for intron removal or cleavage
2. N_en_: Nascent RNAs that have passed the 3’SS but not the cleavage site, which are eligible for intron removal but not cleavage
3. N_rn_: Nascent RNAs that have been spliced but have not passed the cleavage site, which are not eligible for cleavage
4. N_ee_: Nascent RNAs that have passed the cleavage site but have not been spliced, which are eligible for both splicing and cleavage.
5. N_re_: Nascent RNAs that have been spliced and have passed the cleavage site, which are eligible for cleavage.

Let *λ*_*s*_ > 0 and *λ*_*C*_ > 0 be the splicing rate and the cleavage rate, respectively, and *τ*_*s*_ and *τ*_*C*_ be the time it takes to transcribe from the transcription start site to the 3’SS and the cleavage site, respectively, and 0 < *τ*_*s*_ < *τ*_*C*_. Let *dr*(*t*) be the initiation rate of nascent RNA at time *t*, and *dr*(*t*) = 0 for *t* ≤ 0. The change in the nascent RNAs in state N_xy_ at time *t* can be described using the following equations.

The change of nascent RNAs in state N_nn_ at time *t* can be written as:

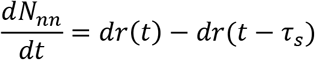

where the right-hand side is the amount of nascent RNAs entering this state, *dr*(*t*), subtracted by the number of nascent RNAs that are leaving the state by passing the SS, *dr*(*t* − *τ*_*s*_).

The change in nascent RNAs in the state N_en_ at time *t* can be written as:

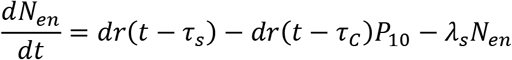

where the right-hand side is the amount of nascent RNAs are entering the state by passing the SS, *dr*(*t* − *τ*_*s*_), subtracted by the nascent RNAs that are passing the cleavage site without splicing, *dr*(*t* − *τ*_*C*_)*P*_10_, and nascent RNAs that are leaving the state via splicing *λ*_*s*_*N*_*en*_. Note that P_10_ is the probability that a nascent RNA reached the cleavage site without being spliced (see below).

The change of nascent RNAs in state N_rn_ at time *t* can be written as:

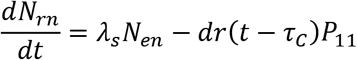

where the right-hand side is the amount of nascent RNAs that are being spliced *λ*_*s*_*N*_*en*_, subtracted by the amount that are passing the cleavage site already spliced, *dr*(*t* − *τ*_*C*_)*P*_11_. Note that P_11_ is the probability that a nascent RNA reached cleavage sites already spliced (see below).

The change of nascent RNAs in state N_ee_ at time *t* can be written as:

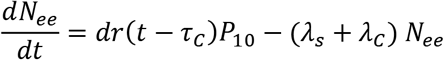

where the right-hand side is the amount of nascent RNAs that are passing the cleavage site without having been spliced, *dr*(*t* − *τ*_*C*_)*P*_10_, subtracted by the amount of nascent RNA that are being spliced or cleaved, (*λ*_*s*_ + *λ*_*c*_) *N*_*ee*_.

Finally, the change of nascent RNAs in state N_re_ at time *t* can be written as:

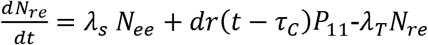

where the right-hand side is the amount of nascent RNAs that are being spliced, added those that are passing the cleavage site already spliced, subtracted by those that are being cleaved.

Based on the exponential distribution, the probabilities P_10_ and P_11_ are as follows:

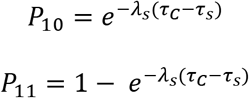

Nascent RNA formed after DRB release is modeled as:

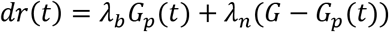

where *G* is the total amount of that gene present in the assay, *λ*_*b*_ is rate of DRB release/pTEF-b reactivation and 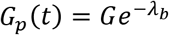 is the amount of nascent RNA that was initiated right after DRB release; *λ*_*n*_ is the initiation rate of nascent RNA; and *λ*_*n*_(*G* − *G*_*p*_(*t*)) is the amount of nascent RNA that were re-initiated.

The above equations were solved sequentially, and we were able to obtain closed formed solutions manually. More complicated event types involve more nascent RNA states (as shown in Fig. S1A), and they can be described similarly using differential equations as illustrated above, and although it can get quite complicated, closed form solutions have been obtained manually for each.

#### mRNA contamination

While the poly(A) depletion described above removes the bulk of mRNA contamination, residual mRNA in the nascent RNA purification would affect the estimations of splicing kinetics without being taken into consideration computationally. Assuming that the mRNA contamination from Drosophila RNA is proportional to the mRNA contamination from the human samples to which it was spiked in, we can use the spiked-in Drosophila RNA to obtain contamination factors:

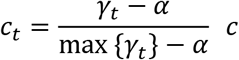

where both *c* and α are estimated together with parameters of gene reactivation, spawn, elongation, splicing and cleavage rates during the optimization process (see below), and *γ*_*t*_ are the observed exon/intron ratio in sequenced Drosophila RNAs from time t, that is, the total number of reads mapped to Drosophila exon over the total number of reads mapped to intron. See Table 3 for the *γ*_*t*_ value used for each time point within each biological replicate. In the above formula, *c* is the reference contamination level for one of the time points; without loss of generality, we let it be of the time point which has the greatest contamination. And 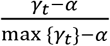 is the ratio between other time point and the reference time point. Note that the sequencing reads come from three categories, exons in nascent RNAs, not yet spliced introns in nascent RNAs, and mRNA contamination, which are also primarily from the exonic regions. Therefore, the observed Drosophila RNAs can be written as *C*_*t*_*e*_*D*_ + *e*_*D*_ + *i*_*D*_, where *e*_*D*_ is the exons, and *i*_*D*_ is the introns that are not spliced, and the exon to intron ratio among the observed RNA is *γ*_*t*_ = (*C*_*t*_ + 1)*α*, where 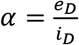 represents unknown exon/intron ratio among nascent RNAs in Drosophila cells, which is stable at different time-points since they are derived from the same pool of Drosophila cells. Since *γ*_*t*_ − *α* = *αC*_*t*_ and the greatest *γ*_*t*_ corresponds to the time point with the greatest contamination, we have *C*_*t*_ /*C* = (*γ*_*t*_ − *α*)/(max {*γ*_*t*_} − *α*), hence the formulae for the contamination factors. Furthermore, in theory, *α* is greater than the exon/intron ratio in the Drosophila genome, which is roughly 0.31 and is less than any observed exon/intron ratios observed across the SKaTER-seq time points. This provides the lower and upbound of *α* in our optimization procedure (see below).

**Table 3.** *γ*_*t*_ for estimation of contamination.

|  | 0min | 5min | 10min | 15min | 20min | 25min | 30min | 35min | SS |
| --- | --- | --- | --- | --- | --- | --- | --- | --- | --- |
| IMR90 replicate 1 | 0.6853 | 0.9921 | 0.6956 | 0.7152 | 0.7448 | 0.8449 | 1.0165 | 0.8292 | 0.5873 |
| IMR90 replicate 2 | 1.0524 | 1.1067 | 0.7980 | 0.8566 | 0.8146 | 0.8164 | 0.8526 | 0.8780 | 1.0951 |
| A549 replicate 1 | 1.3033 | 1.2437 | 1.1685 | 1.0364 | 0.9228 | 1.0940 | 1.1680 | 1.1259 | 1.0271 |
| A549 replicate 2 | 0.7578 | 0.6959 | 0.6546 | 0.6097 | 0.5882 | 0.7418 | 0.7061 | 0.7093 | 0.6657 |

Since the parameters involved in the estimation of mRNA contamination are themselves estimated during the optimization procedure (see below), they are not simply subtracted from the actual gene coverage data prior to optimization. Instead, the contamination factors are used in our model to add the effect of mRNA contamination onto the predicted coverage. The contamination factor represents the fraction of a predicted pool of mRNA produced over between 5 and 9 hours following DRB washout (a time that approaches steady state transcription) that is then added to the predicted nascent coverage to give the overall predicted coverage across the gene.

#### Differential optimization algorithm

A differential evolution optimization algorithm^16,17^ was used along with the model of nascent RNA transcription and co-transcriptional processing described above to determine the set of rates that best fit the SKaTER-seq data. We start with a population of potential solutions, each of which contains numerical values for all of the parameters (rates) needed to obtain the predicted coverage for an individual gene. Rates making up each solution in the first population are randomly chosen between the set upper and lower bounds for that rate (Table 1) from a uniform distribution. For each solution, we model the coverage based on the rates from the potential solution, then the residual sum of squares (comparing the modeled coverage and the SKaTER-seq coverage across time points) was used to determine the “fitness” of each potential solution. Better solutions will have a smaller RSS indicating that the predicted coverage closely matches the observed coverage. Each parameter of the potential solution is simultaneously optimized until all rates converged. We determine if rates have converged by comparing the maximum RSS calculated for a solution in a generation and the minimum RSS calculated for a solution. If this metric drops below 0.1, the solution has converged.

### SKaTER-seq rate analysis

#### Filtering rates

Rates produced by the SKaTER-seq pipeline were further filtered based on the coverage of the gene in the nascent RNA-seq and the stability of the rate over multiple optimizer runs. A BED file of all genes based on the transcription start sites and stops sites in RefSeq was created and the number of nascent RNA-seq reads per gene was computed using the multicov tool in the BEDtools suite (v2.29.2^42^). Coverage was calculated by dividing the number of reads per gene by the length of the gene (kb). To be included in further analysis, we required that each gene have a coverage of at least 5 paired reads/kb.

To evaluate the stability of the rates over multiple runs, we calculated the variance of the log_2_(rate) over at least 3 runs of the optimizer. To be included in further analysis, we required the variance to be below 0.1. We also required that the rate was not within 5% of the set bounds for that rate type (therefore removing rates that have “maxed out” or “bottomed out” during optimization, see Table 1 for bounds and Fig. S2A for the number of expressed genes with solved rates).

Spawn rates are relative within a cell type and cannot be directly compared across cell types due to the amplification step in library preparation for high-throughput sequencing (as they are dependent on the number of loci being sequenced). Unlike spawn rates, elongation, splicing and cleavage rates are absolute and so can be compared across conditions.

#### GC content

To determine GC content within specific loci, BED files were created where each line was a region to be analyzed. The sequences and fraction GC content were determined using the nucBed tool in the BEDtools suit (v2.29.2^42^).

#### PSI prediction

Using our model of nascent RNA biogenesis and processing (described above), we simulated the steady-state pool of mRNA at 24 hours post-DRB release using the rates generated from the optimization algorithm, assuming all transcripts have similar decay rates. The predicted PSI for each alternative splicing event was calculated as (# transcripts containing the alternative exon / total # of transcripts). This metric was compared to the PSI calculated from total RNA-seq data as described above.

#### Hi-C analysis

Paired-end FASTQ sequences were first evaluated as separate, single-end sequencing files. Sequences were trimmed at the restriction sites using homerTools^26^ then aligned to the human genome (hg19) using botwie2 (v2.3.3.1^43^). Paired-end tag directories were made using the HOMER makeTagDirectory program, combining the sequences from all biological replicates performed for each cell line. Then hic files were created using the HOMER tagDir2hicFile program. Full matrices were extracted from the hic files using the Juicer Tools^27^ dump command at a resolution of 50 kb. TADs were called using the Juicer Tools matrices and the domain calling software program published by Dixon et al^24^. AB compartments were determined at a resolution of 1 Mb using the Eigenvector algorithm on the Juicer platform^27^.

#### Permutation tests

##### Variance of splicing rates within genes (Fig. 2E-G)

For each gene, splicing rates of constitutive introns that passed the filtering steps listed above in both biological replicates were collected, and the observed total variance was calculated:

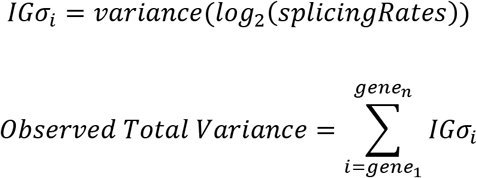

Where *IGσ*_*i*_ is the intra-gene variance of the log_2_(rate). In order to be included in the analysis, the gene must contain at least 3 constitutive introns with stable rates. The observed total variance is the sum of the *IGσ*_*i*_ across all genes. The closer the observed total variance is to zero, the more coordinated rates are within genes. The rates of individual introns were then shuffled across all genes included in the analysis, and the total variance was recalculated. This randomization and recalculation was performed 100,000 times and the number of times that the total variance based on permuted rates was less than the observed total variance based on SKaTER-seq rates was counted. The observed/expected metric was calculated, and significance was determined using the p-value of the z-score comparing the observed total variance to the expected distribution.

##### Variance of rates within windows/genomic compartments (Extended Data Fig. 4B-D, Fig. 3C)

The intronic splicing rates or gene-average rates in a window were used (for gene-average splicing rates, each gene must have at least 3 confidently called splicing rates to be averaged).

Then the window-level total variance was calculated:

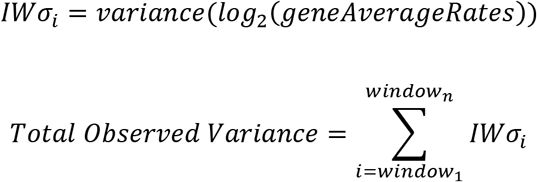

where *IWσ*_*i*_ is the intra-window variance of the log_2_(gene-average rates). In order to be included in the analysis, the window must contain 3 genes with calculated gene-average rates. The variance was summed across all windows to get the total observed variance. The permutation test was performed as described above.

##### Sum of delta PSI values within windows (Extended Data Fig. 6A, C, Fig. 4E)

Delta PSI (ΔPSI) values were computed using rMATS (v3.2.5^37^). Positive ΔPSI values indicate more exon skipping in the second condition when compared to the first, while negative ΔPSI values indicate more exon inclusion in the second condition. Events were included in the analysis if the ΔPSI was greater than or equal to 0.05, and if the variance of the PSI values within the replicates of each condition was less than 0.1. Furthermore, if the genomic position of multiple identified events overlapped, only the event with the highest expression (based on total junctional read counts across each replicate) was considered. Windows were included in the analysis if they contained at least two alternative splicing events. The ΔPSI values were converted to +1 or −1 depending on their sign to get the ordinal ΔPSI, then the window level total variance was calculated as follows:

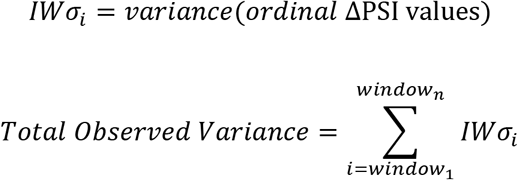

The ΔPSI values were then shuffled across all events included in the original analysis and the total ordinal ΔPSI variance was recalculated. This randomization and recalculation was performed 100,000 times and the number of times that the observed total ordinal ΔPSI variance was less than the expected total ordinal ΔPSI variance was counted. The empirical p value was calculated to determine if the coordination of ΔPSI in the actual data was significantly greater than that of the randomly distributed data. The observed/expected metric was calculated and the significance was determined based on the p-value of the z-score comparing the observed ordinal ΔPSI variance to the expected distribution.

#### Unsupervised genomic segmentation (Fig. 3E, 4F)

To segment the genome based on gene-average rates or ΔPSI values, we used the binary segmentation change point algorithm in the Python package Ruptures^44^. The segments encompass genes with solved rates, therefore the start and stop positions of the segments are definitionally at gene start or stop positions. The “true” change points are still ambiguous, falling somewhere between segments, within the change point zones (CPZ). In order to determine how much shared information exists between two sets of segmentation (i.e. unsupervised segments based on SKaTER-seq rates or ΔPSI values and established segments such as TADs or A/B compartments), we calculated the variation of information metric (VOI) for all pairwise combinations of segments^45^.

If a pair of segments overlap perfectly, the VOI will be 0; the VOI will increase if segments have less or no overlap). The VOI is summed over all pairwise comparisons to get the observed VOI. The location of the change point zones is then shuffled and the total VOI is recalculated. This randomization and recalculation was performed 100,000 times and the number of times that the observed total VOI was less than the expected total VOI was counted. The empirical p value was calculated to determine if the change point zones based on the segmentation of the actual data are more similar to the established segments than would be expected by random chance. The observed/expected metric was calculated and the significance was determined based on the p-value of the z-score comparing the observed total VOI to the expected distribution.

